# The common chromosomal periodicity of transcriptomes is correlated with the bacterial growth rate

**DOI:** 10.1101/2020.05.24.113886

**Authors:** Motoki Nagai, Masaomi Kurokawa, Bei-Wen Ying

**Affiliations:** Graduate School of Life and Environmental Sciences, University of Tsukuba, 1-1-1 Tennoudai, Tsukuba, Ibaraki 305-8572, Japan

**Author notes:** Correspondence (BWY).

**Keywords:** chromosomal periodicity, transcriptome, growth rate, population fitness

## Abstract

The growth rate, representing the fitness of a bacterial population, is determined by the whole transcriptome. Chromosomal periodicity is a representative overall feature of the whole transcriptome; however, whether and how it is associated with the bacterial growth rate are unknown. To address these questions, we analyzed a total of 213 transcriptomes of genetically differentiated *Escherichia coli* strains growing in an assortment of culture conditions varying in terms of temperature, nutrition level and osmotic pressure. Intriguingly, the Fourier transform identified a common chromosomal periodicity of transcriptomes, which was independent of the variation in genomes and environments. In addition, fitting of the theoretical model found that the amplitudes of the periodic transcriptomes were significantly correlated with the growth rates. This novel finding successfully identified a single parameter representing the global pattern of the whole transcriptome for the first time and indicated that bacterial growth was correlated with the magnitude of chromosomal differentiation in gene expression. These results provided an alternative global parameter for evaluating the adaptiveness of a growing bacterial population and provided a quantitative rule that makes it possible to predict the growth dynamics according to the gene expression pattern.

## Introduction

The growth rate in the exponentially growing phase is the most important parameter representing both genetic and environmental influences on bacterial growth dynamics. Predicting the growth rate of a growing bacterial population according to the intrinsic status and/or extrinsic conditions is highly desirable. To date, extensive studies involving the systematic quantitative investigation of bacterial growth have been performed. By using systematic genetic constructs, e.g., single-gene knockout ^1^ and genome reduction, ^2,3^, the contributions of both single genes and large genomic fragments to bacterial growth were quantitatively evaluated ^4-6^. The finding of a correlation between the genome and the growth rate strongly suggests that population fitness is linked to genome-wide features in vivo ^5,7^.

The transcriptome, which illustrates a global view of the transcriptional abundance of all the genes distributed in the genome, is reorganized constantly in response to genomic and environmental perturbations ^8-10^. As the transcriptome is known to be associated with population fitness ^11-13^, the contribution of the transcriptome to population fitness is of great interest. Our previous studies reported the coordination of gene expression with the growth rate ^13^ and the linkage between transcriptome reorganization and increases in fitness in adaptation and evolution ^14,15^. These findings indicated that the whole transcriptome, rather than the specific regulation of limited gene groups, increased population fitness. Whether and how the whole transcriptome is linked to population fitness remains unknown.

A single parameter representing the whole transcriptome is critical for determining the linkage if it exists. Previous studies successfully demonstrated that the power law (Zipf’s rule) was a universal principle governing the transcriptome in living organisms ^16,17^; however, we failed to find the linkage between this law and the growth rate ^18^. As an alternative global feature representing the transcriptome, chromosomal periodicity has been proposed ^19,20^, which is determined using the Fourier transform, a mathematical method used to estimate the periodic patterns in an entire dataset according to the sinusoidal wave ^21^. Computational analyses identified some particular periods associated with bacterial transcriptomes ^19,20^, which were supported by the molecular functions and/or mechanisms related to the chromosomal topology ^22-24^. These findings of chromosomal periodicity were used to obtain static snapshots of the whole transcriptome, and whether and how the chromosomal periodicity of the transcriptome is linked to the growth rate are under investigation.

In the present study, a total of 213 growth profile-associated transcriptomes were analyzed that represent an assortment of genetically differentiated *E. coli* cells growing under or responding to various environments. This study seeks to determine whether the chromosomal periodicity of the transcriptome is robust or plastic in response to environmental and genetic differentiation and whether and how chromosomal periodicity is coordinated with bacterial growth.

## Materials and Methods

### E. coli strains and growth conditions

Three types of *E. coli* genomes were included in the transcriptome analyses: the full-length genomes of MG1655 and DH1 and the partial genome of MDS42 ^2^. A number of genetically engineered strains were comprised of genomes of types DH1 ^14,25^ and MDS42 ^13^, which led to a total of 20 different genomic backgrounds. The growth media were all based on the minimal medium M63 ^26^; if required, the medium was supplemented with factors, including amino acids, to compensate for the interruption of gene function resulting from genetic engineering. In addition, the growth temperature was also varied. Nine different media and seven different temperatures were used, which resulted in a total of 16 different environmental conditions. The *E. coli* growth status was evaluated in the exponential growth phase and the stress response phase. Only the transcriptomes associated with the exponential growth phase were linked with precise growth rates (165 transcriptomes) and were subjected to correlation analysis. The details of the genomic backgrounds and the environmental conditions can be found in previous reports ^13-15,18,25,27,28^.

### Transcriptomes

The transcriptomes used in the present study were acquired from microarray raw data sets assigned the GEO access numbers of GSE33212, GSE49296, GSE55719, GSE52770 and GSE61749 by using the customized platform EcFS ^29^. The finite hybridization model ^30^ was applied to determine the gene expression levels, which were calculated as the log-scale mRNA concentrations (pM). Data filtering, normalization and averaging of the biological replicates for the subsequent transcriptome analyses were described previously. The resulting transcriptomes were associated with the growth profiles, genomic backgrounds and environmental conditions. The details were previously described in the corresponding studies ^13-15,25,27^. A total of 213 transcriptomes, comprising 72 combinations (biological repetition, N=2∼7) that varied in terms of the genomic background and environmental conditions as described above, were included in the analyses.

### Computational analyses

All computational analyses were performed with R ^31^. The gene expression levels on a logarithmic scale were used for the analyses as described previously ^18,32^. A total of 165 transcriptomes representing the exponential growth phase associated with the repeated growth assay were subjected to correlation analysis of the growth rate (**r**) and the periodic parameters (**a, b** and **c**). This resulted in 42 combinations of genomic and environmental variations. The statistical significance of the Pearson correlation coefficients was evaluated by the t-test. The Z-score was used for the standardization of the MG1655 transcriptomes obtained from the present data sets and the GyrA Chip-seq data for MG1655 obtained from another study ^20^. The Z-scores of both the gene expression and the GyrA binding activity were calculated and were averaged with a sliding distance of 1 kb for the correlation analysis and the Fourier transform.

### Evaluation of chromosomal periodicity

A standard Fourier transform was employed to determine the chromosomal periodicity of the transcriptomes and the GyrA binding activity by using the periodogram function in R. All 213 transcriptomes were subjected to the Fourier transform, in which the expression data for 4393, 3760 and 4377 genes in the *E. coli* strains with the genomic backgrounds of MG1655, MDS42 and DH1 were used, respectively. The CDS information for MG1655, MDS42 and DH1 were obtained from the DDBJ databanks under the accession IDs U00096, AP012306 and AP012030, respectively. The sizes of the genomes used for the Fourier transform were 4642, 3976 and 4622 kb and corresponded to the genomic backgrounds of MG1655, MDS42 and DH1, respectively. The chromosomal periodicities of both the transcriptomes and the GyrA binding activity were evaluated with a sliding distance of 1 kb and are shown in 100-kb bins. The approximate curves of the periodicity were calculated using the highest peak of the periodogram and were fitted by minimizing the square error of the approximate curve and the series of expression values. The statistical significance of the periodicity was assessed with Fisher’s g test ^21^, which was performed using the GeneCycle package in R.

## Results and Discussion

### The common chromosomal periodicity of the transcriptome

To investigate whether the growth conditions and the genomic background influenced the chromosomal periodicity of the transcriptome, a total of 213 *E. coli* transcriptome data sets, which were associated with the growth profiles and were acquired with the same microarray platform, were used in the present study. The growth conditions were varied in terms of temperature, nutrition level and osmotic pressure, and there was a large variation in the genomic backgrounds (as described in the Materials and Methods). Chromosomal fluctuations in gene expression were confirmed (Fig. 1A), and chromosomal periodicity was evaluated with the Fourier transform.

**Figure 1.**
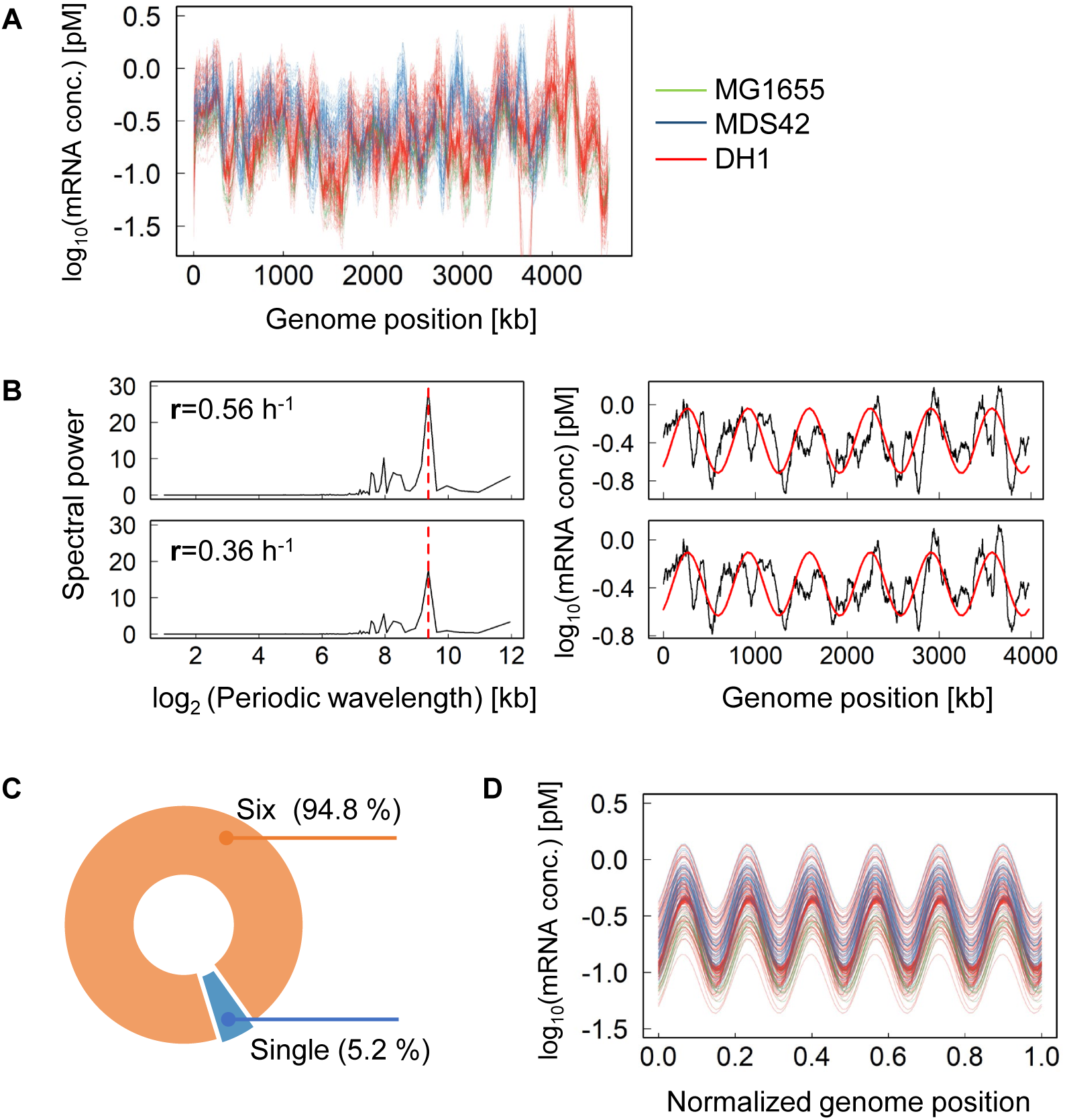
Chromosomal periodicity of the transcriptome. **A**. All transcriptomes used in the present study. The transcriptional levels of every 1 kb sliding window and 100 kb smoothing are shown. The color variation indicates the transcriptomes of individual strains. **B**. Periodograms of the transcriptomes. Two transcriptomes with different growth rates are shown. The growth rates, **r**, are indicated in the insets. The left and right panels represent the distributions of the Fourier-transformed periodic wavelengths on a logarithmic scale and the estimated chromosomal periodicity of the transcriptome, respectively. The broken lines and solid curves in red indicate the highest power spectra (the max-peak) estimated by the Fourier transform and the corresponding fitted period of the transcriptome, respectively. The transcriptional levels for every 1 kb sliding window and 100 kb smoothing are shown. **C**. Distribution of the periods corresponding to the max-peak. Orange and blue indicate the ratios of the six periods and the single period among the 213 total transcriptomes, respectively. **D**. Overlapping periods of the transcriptomes. The chromosomal periodicity of 202 transcriptomes showing six periods (orange in **C**) were plotted together. The color variation corresponds to that shown in **A**.

Intriguingly, the analysis results showed the high level of common chromosomal periodicity of the transcriptomes, which was independent of the growth conditions and the genomic backgrounds. For instance, the most significant spectral powers (*i*.*e*., the max peak) identified in two transcriptomes associated with different growth rates (**r**) were exactly the same (Fig. 1B, left panels). Consequently, this resulted in an identical chromosomal periodicity (Fig. 1B, right panels), although the two transcriptomes represented the *E. coli* cells growing at different temperatures. Overall, 202 out of 213 transcriptomes presented a universal chromosomal periodicity of six periods (Fig. 1C) as the highest priority in the Fourier transform. Of note, all 11 exceptions showed a chromosomal periodicity of six periods as the second priority. As the statistical significance of the chromosomal periodicity was further proven by Fisher’s g test for all transcriptomes (Fig. S1), the determination of the common chromosomal periodicity of transcriptomes, which consisted of six periods, was highly reliable. This result agreed with those of previous studies reporting periodic transcriptomes either in wild-type *E. coli* strains or under regular growth conditions ^20,27,33^.

In addition, the periodicity of the chromosomal dynamics of the transcriptomes was somehow synchronized. Despite the large variation in both the genomic backgrounds of the *E. coli* strains and the environmental conditions of the population growth, the six periods of a total of 202 transcriptomes almost overlapped (Fig. 1D). The similar directional changes in gene expression among the genomic positions further demonstrated the universality of the chromosomal periodicity of the transcriptomes irrespective of genetic and environmental disturbances. This was the first finding that revealed that neither the number nor the wavelength of the periods was linked to bacterial growth.

### Correlation between the growth rate and the amplitude of the periodic transcriptome

Whether there was any parameter representative of the periodic transcriptomes linked to bacterial growth was further investigated. The gene expression level, *Exp*(*x*), was related to the genome position (*x*) of the corresponding gene. The parameters affecting the chromosomal periodicity of the transcriptome were theoretically defined in the following formula (Eq. 1).

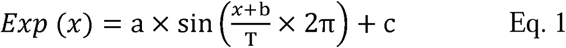

Here, the parameters **a, b**, and **c** represented the amplitude of the period, the phase of the period (i.e., the genomic position of the period initiation), and the mean transcriptional level, respectively (Fig. 2A). The estimation of the three parameters was performed by minimizing the square error in the curve fitting. The constant T was the wavelength of the period of the highest spectral power estimated by the Fourier transform. A total of 165 transcriptomes, which represented the exponential growth phase and were associated with highly precise growth rates (**r**), were subjected to theoretical fitting with Eq. 1.

**Figure 2.**
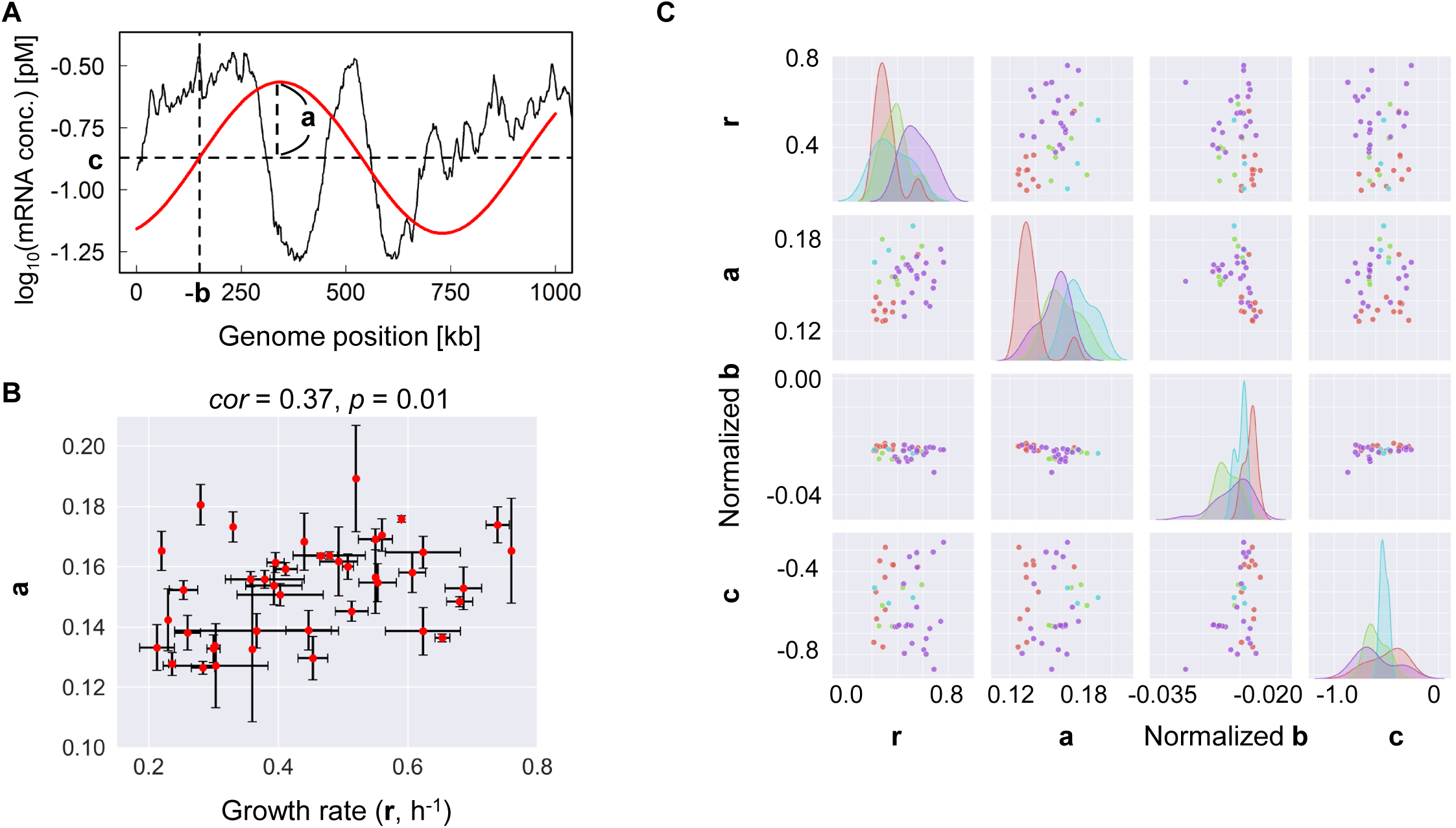
Correlation between the chromosomal periodicity and the growth rate. **A**. Illustration of the parameters defined for the periodic transcriptome. The parameters (**a, b** and **c**) used in Eq. 1 are indicated. Black and red lines represent the transcriptome and the fitted period, respectively. **B**. Scatter plot of the amplitude of the six periods and the growth rate. The standard errors of the biological replicates are indicated. The correlation coefficient and its significance are indicated. **C**. Relationships among the parameters defined for the chromosomal periodicity and the growth rate. The relationship between any two of **a, b, c** and **r** is shown in matrix form. Pink, blue, green and purple represent the environmental variations in temperature, osmotic pressure, and nutritional level and normal conditions, respectively.

The theoretical fitting successfully identified a significant correlation between the growth rate and the amplitude of the periodic transcriptome. The values of **a, b** and **c** calculated by curve fitting were averaged among the biological repeats, which led to 42 combinations that varied in terms of the genomic background and/or environmental conditions. Note that parameter **b** was further normalized because of the variation in the genome length. The parameter **a** was positively correlated with the growth rate (Fig. 2B), whereas such a correlation was not detected for the parameters **b** and **c** (Fig. S2). The analysis clearly determined a simple correlation between the growth rate and the amplitude of the period, although the 165 transcriptomes with 42 combinations largely differed in terms of the genotypes and environments. This strongly suggested that population fitness was correlated with the magnitude of differential transcription along the chromosome.

Further investigations of the contributions of the genomic background and the environmental conditions failed to observe any significant relationship with the growth rate. According to previous reports ^13-15,18,25^, four types of environmental variations were roughly categorized as normal conditions and conditions with changes in temperature, nutrition level and osmotic pressure. The distributions of the four parameters representing the growth and the periodicity of the transcriptome were largely dissimilar among the four categories (Fig. 2C), which reflected the properties of the data sets. No environmentally dependent feature or correlation among the parameters **a, b**, and **c** was found (Fig. 2C). Additionally, genetic engineering might affect the phase of the period (*i*.*e*., normalized **b**), as a difference was detected between the wild-type genome and the other genomes (Fig. S3). The genetic reconstruction possibly interrupted the genomic position of the period initiation, although more datasets were required to support this assumption.

### Mechanisms of the universal chromosomal periodicity of transcriptomes

To understand the universality of the periodicity of transcriptomes, a simple assumption was made that the essential genes determined the six periods. However, neither deleting the essential genes from the transcriptome data nor substituting the true expression values with zero altered the common periodicity of the transcriptome (Fig. S4). This result indicated that the periodicity of the transcriptome was not simply due to the genomic localization of the essential genes, although it is unclear whether and how the absence of the essential genes triggered the transcriptional changes of the nonessential genes, as deleting the essential genes from the genome was impractical.

As the chromosomal structure might contribute to transcriptional activity ^32,34-36^, whether the common chromosomal periodicity of the transcriptome was attributed to the chromosomal organization was determined. The macrodomain model was proposed for the *E. coli* chromosome, with four structured domains and two nonstructural regions ^37-40^. The normalized periodicity of the transcriptomes showed that the six periods were roughly positioned within the six domain regions of the *E. coli* chromosome (Fig. 3A), which was consistent with previous findings ^20,27,33^. The highly overlapping phases of the periodic transcriptome (Fig. 3A) suggested that the chromosomal macrodomain structure was robust against genomic and environmental disturbances.

**Figure 3.**
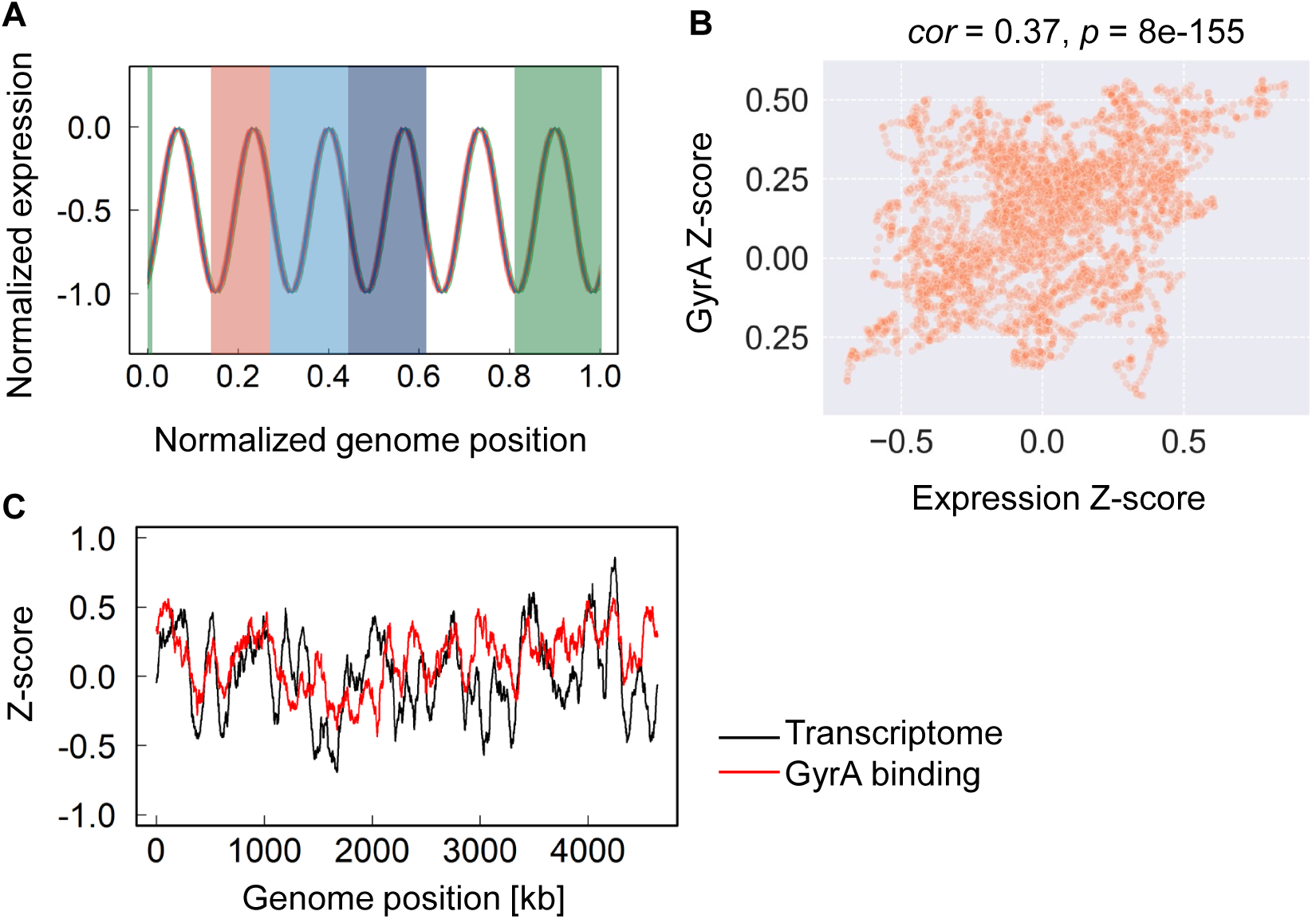
Comparison of the common periodicity of the transcriptomes to chromosome structures. **A**. Relationship between the chromosomal macrodomains and the periodic transcriptomes. The normalized periodic transcriptomes are shown. Four macrodomain regions and two nonstructural regions are shown in solid color and as transparent, respectively. The macrodomains of the Ori, Right, Ter and Left regions are shown in green, red, light and dark blue, respectively. **B**. Scatter plot of the transcriptome versus GyrA binding activity in MG1655. Standardization of both the mean expression levels and the GyrA Chip-seq data were performed by determining the z-score. The correlation coefficients and the statistical significance are indicated. **C**. Comparison of the transcriptome and the GyrA binding activity of the wild-type genome MG1655. Red and black curves indicate GyrA binding activity and the transcriptome, respectively. Both were calculated using a 1 kb sliding window and are shown according to the 100-kb moving average.

Moreover, the molecular mechanism related to the DNA topology probably played a role in determining the chromosomal periodicity of the transcriptome. Bacterial chromosomal structures are highly dynamic and compacted in association with nucleoid-associated proteins (NAPs). Previous studies indicated that the chromosomal supercoiling of domains ∼10 kb size ^20,22^ was potentially attributed to the chromosomal localization of nucleoid-associated proteins, *e*.*g*., H-NS ^41^, and that of domain 600∼800 kb size might be triggered by DNA gyrase ^20^. As the common period identified in the present study was ∼700 kb, the participation of the subunit of the DNA gyrase, GyrA, was confirmed. A highly significant correlation was verified between the transcriptome of the wild-type strain MG1655 in the present study and the abundance of chromosomally bound GyrA in a previous report ^20^ (Fig. 3B). Such a correlation seemed to be common in all transcriptomes (Fig. S5) if the properties of the binding of GyrA to the genome remained unchanged. The correlation between gene expression and binding activity suggested the similarity of the chromosomal periodicity of GyrA binding and that of the transcriptome. Although the six periods were not the first priority for GyrA binding (Fig. S6), the chromosomal periodicities of GyrA binding activity and the transcriptome in MG1655 were somehow coordinated (Fig. 3C).

### Linking the periodicity of the transcriptome to population fitness

The present study successfully found a direct linkage between population dynamics and the whole transcriptome (Fig. 4). The whole transcriptome is influenced by both the genomic background and the environmental conditions; consequently, it determines bacterial growth. Previous studies of transcriptomes successfully classified the genes into diverse categories that functioned either specifically in response to environmental changes or generally in relation to the growth rate ^11-13^. However, whether there was any quantitative relationship directly linking the two global features of growth and the transcriptome remained unaddressed.

**Figure 4.**
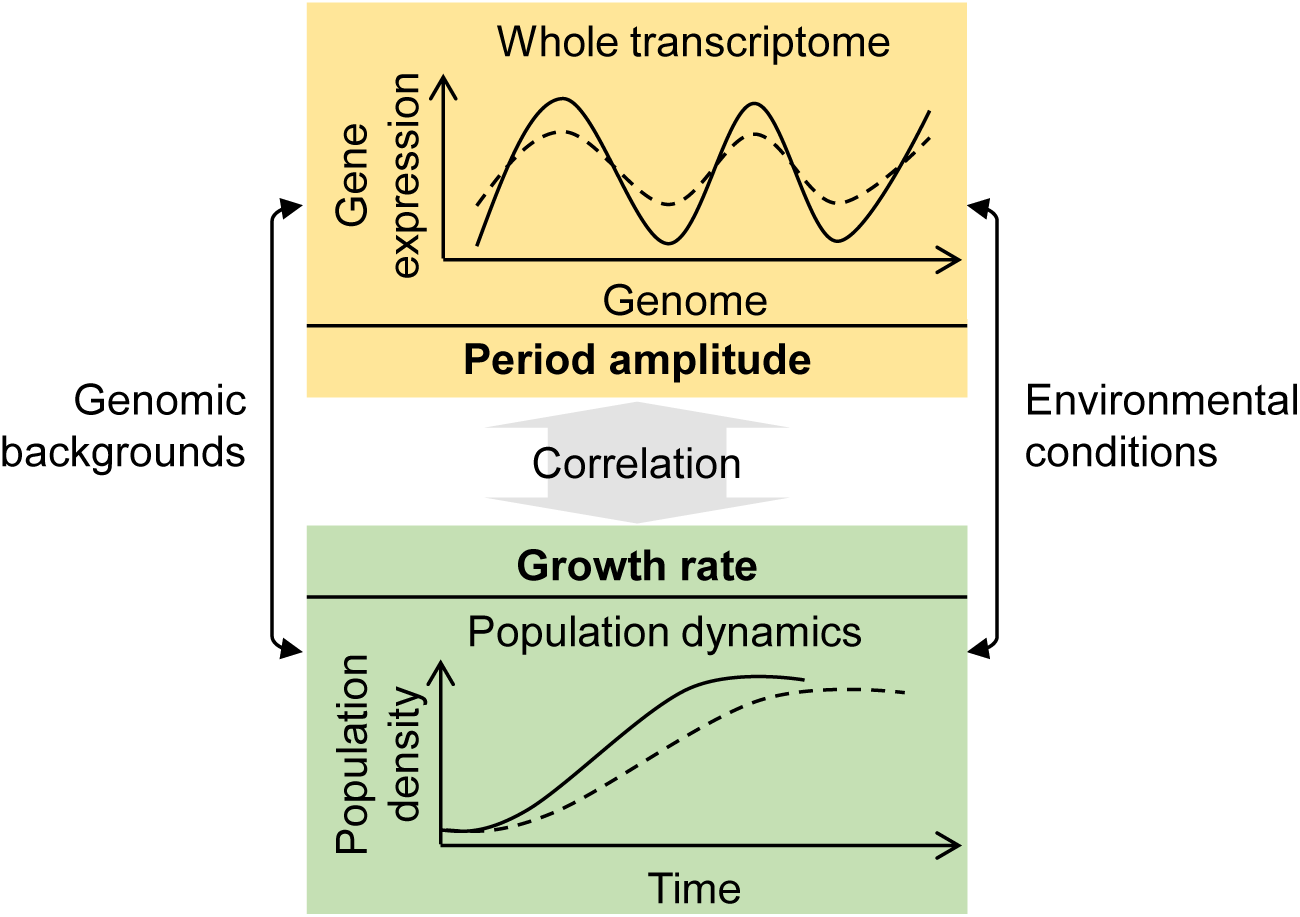
Scheme of the correlation between the growth rate and the chromosomal periodicity of the transcriptome. Shadowed boxes indicate the profiles of the whole transcriptome and the corresponding population dynamics. The broken and solid curves in the upper box indicate the small and large amplitudes of the periodic transcriptomes, respectively, and those in the bottom box indicate slow and fast growth, respectively.

The present study first identified a single parameter that represented well the global features of the whole transcriptome. That is, the amplitude of the chromosomal periodicity of the transcriptome represents the magnitude of the chromosomal differentiation of gene expression. Moreover, fast growth was linked to a large amplitude of the periodic transcriptome (Fig. 4, solid curves). The correlation between the growth rates and the amplitudes of periodic transcriptomes was independent of the environmental conditions and the genomic backgrounds. This novel finding was a breakthrough for understanding how the whole transcriptome determined population fitness because it was the first demonstration that the magnitude of chromosomal differentiation of gene expression was correlated with the growth rate.

The growth rate was the only global parameter representing the adaptiveness of a growing bacterial population. The amplitude of the periodic transcriptome could be considered an alternative global parameter for evaluating population fitness. In addition to network reconstruction ^42^, the assessment of the global pattern of the whole transcriptome might be applied for predicting population fitness, which would be beneficial for industrial applications to substrate production and the fundamental investigation of living principles.

## Acknowledgments

We thank Shigeto Seno for the programming assistance. This work was supported by the JSPS KAKENHI Grant-in-Aid for Scientific Research (B) (grant number 19H03215 (to BWY)).

## References

1. Baba, T., Ara, T., Hasegawa, M., et al. 2006, Construction of Escherichia coli K-12 in-frame, single-gene knockout mutants: the Keio collection. Molecular systems biology, 2, 2006 0008.

2. Posfai, G., Plunkett, G., 3rd, Feher, T., et al. 2006, Emergent properties of reduced-genome Escherichia coli. Science, 312, 1044–1046.

3. Kato, J. and Hashimoto, M. 2007, Construction of consecutive deletions of the Escherichia coli chromosome. Molecular systems biology, 3, 132.

4. Karcagi, I., Draskovits, G., Umenhoffer, K., et al. 2016, Indispensability of Horizontally Transferred Genes and Its Impact on Bacterial Genome Streamlining. Molecular biology and evolution, 33, 1257–1269.

5. Kurokawa, M., Seno, S., Matsuda, H. and Ying, B. W. 2016, Correlation between genome reduction and bacterial growth. DNA research: an international journal for rapid publication of reports on genes and genomes, 23, 517–525.

6. Campos, M., Govers, S. K., Irnov, I., Dobihal, G. S., Cornet, F. and Jacobs-Wagner, C. 2018, Genomewide phenotypic analysis of growth, cell morphogenesis, and cell cycle events in Escherichia coli. Molecular systems biology, 14, e7573.

7. Nishimura, I., Kurokawa, M., Liu, L. and Ying, B. W. 2017, Coordinated changes in mutation and growth rates induced by genome reduction. mBio, 8.

8. Gibson, G. 2008, The environmental contribution to gene expression profiles. Nature reviews. Genetics, 9, 575–581.

9. Jozefczuk, S., Klie, S., Catchpole, G., et al. 2010, Metabolomic and transcriptomic stress response of Escherichia coli. Molecular systems biology, 6, 364.

10. Feugeas, J. P., Tourret, J., Launay, A., et al. 2016, Links between Transcription, Environmental Adaptation and Gene Variability in Escherichia coli: Correlations between Gene Expression and Gene Variability Reflect Growth Efficiencies. Molecular biology and evolution.

11. Lopez-Maury, L., Marguerat, S. and Bahler, J. 2008, Tuning gene expression to changing environments: from rapid responses to evolutionary adaptation. Nature reviews. Genetics, 9, 583–593.

12. Nahku, R., Valgepea, K., Lahtvee, P. J., et al. 2010, Specific growth rate dependent transcriptome profiling of Escherichia coli K12 MG1655 in accelerostat cultures. Journal of biotechnology, 145, 60–65.

13. Matsumoto, Y., Murakami, Y., Tsuru, S., Ying, B. W. and Yomo, T. 2013, Growth rate-coordinated transcriptome reorganization in bacteria. BMC genomics, 14, 808.

14. Murakami, Y., Matsumoto, Y., Tsuru, S., Ying, B. W. and Yomo, T. 2015, Global coordination in adaptation to gene rewiring. Nucleic acids research, 43, 1304–1316.

15. Ying, B. W., Matsumoto, Y., Kitahara, K., et al. 2015, Bacterial transcriptome reorganization in thermal adaptive evolution. BMC genomics, 16, 802.

16. Furusawa, C. and Kaneko, K. 2003, Zipf’s law in gene expression. Phys Rev Lett, 90, 088102.

17. Ueda, H. R., Hayashi, S., Matsuyama, S., et al. 2004, Universality and flexibility in gene expression from bacteria to human. Proceedings of the National Academy of Sciences of the United States of America, 101, 3765–3769.

18. Ying, B. W. and Yama, K. 2018, Gene Expression Order Attributed to Genome Reduction and the Steady Cellular State in Escherichia coli. Frontiers in microbiology, 9, 2255.

19. Allen, T. E., Herrgard, M. J., Liu, M., et al. 2003, Genome-scale analysis of the uses of the Escherichia coli genome: model-driven analysis of heterogeneous data sets. Journal of bacteriology, 185, 6392–6399.

20. Jeong, K. S., Ahn, J. and Khodursky, A. B. 2004, Spatial patterns of transcriptional activity in the chromosome of Escherichia coli. Genome biology, 5, R86.

21. Wichert, S., Fokianos, K. and Strimmer, K. 2004, Identifying periodically expressed transcripts in microarray time series data. Bioinformatics, 20, 5–20.

22. Postow, L., Hardy, C. D., Arsuaga, J. and Cozzarelli, N. R. 2004, Topological domain structure of the Escherichia coli chromosome. Genes Dev, 18, 1766–1779.

23. Abel, J. and Mrazek, J. 2012, Differences in DNA curvature-related sequence periodicity between prokaryotic chromosomes and phages, and relationship to chromosomal prophage content. BMC genomics, 13, 188.

24. Dorman, C. J. 2013, Genome architecture and global gene regulation in bacteria: making progress towards a unified model? Nature Reviews Microbiology, 11, 349–355.

25. Yama, K., Matsumoto, Y., Murakami, Y., et al. 2015, Functional specialization in regulation and quality control in thermal adaptive evolution. Genes to cells: devoted to molecular & cellular mechanisms.

26. Kurokawa, M. and Ying, B. W. 2017, Precise, High-throughput Analysis of Bacterial Growth. J Vis Exp.

27. Ying, B. W., Seno, S., Kaneko, F., Matsuda, H. and Yomo, T. 2013, Multilevel comparative analysis of the contributions of genome reduction and heat shock to the Escherichia coli transcriptome. BMC genomics, 14, 25.

28. Ying, B. W., Yama, K., Kitahara, K. and Yomo, T. 2016, The Escherichia coli transcriptome linked to growth fitness. Genom Data, 7, 1–3.

29. Furusawa, C., Ono, N., Suzuki, S., Agata, T., Shimizu, H. and Yomo, T. 2009, Model-based analysis of non-specific binding for background correction of high-density oligonucleotide microarrays. Bioinformatics, 25, 36–41.

30. Ono, N., Suzuki, S., Furusawa, C., et al. 2008, An improved physico-chemical model of hybridization on high-density oligonucleotide microarrays. Bioinformatics, 24, 1278–1285.

31. Ihaka, R. and Gentleman, R. 1996, R: A language for data analysis and graphics. Journal of computational and graphical statistics, 5, 299–314.

32. Ying, B. W., Seno, S., Matsuda, H. and Yomo, T. 2017, A simple comparison of the extrinsic noise in gene expression between native and foreign regulations in Escherichia coli. Biochemical and biophysical research communications, 486, 852–857.

33. Allen, T. E., Price, N. D., Joyce, A. R. and Palsson, B. O. 2006, Long-range periodic patterns in microbial genomes indicate significant multi-scale chromosomal organization. PLoS computational biology, 2, e2.

34. Muskhelishvili, G., Forquet, R., Reverchon, S., Meyer, S. and Nasser, W. 2019, Coherent Domains of Transcription Coordinate Gene Expression During Bacterial Growth and Adaptation. Microorganisms, 7, 694.

35. Meyer, S., Reverchon, S., Nasser, W. and Muskhelishvili, G. 2018, Chromosomal organization of transcription: in a nutshell. Curr Genet, 64, 555–565.

36. Sobetzko, P., Travers, A. and Muskhelishvili, G. 2012, Gene order and chromosome dynamics coordinate spatiotemporal gene expression during the bacterial growth cycle. Proceedings of the National Academy of Sciences of the United States of America, 109, E42–50.

37. Espeli, O., Mercier, R. and Boccard, F. 2008, DNA dynamics vary according to macrodomain topography in the E. coli chromosome. Molecular microbiology, 68, 1418–1427.

38. Niki, H., Yamaichi, Y. and Hiraga, S. 2000, Dynamic organization of chromosomal DNA in Escherichia coli. Genes Dev, 14, 212–223.

39. Valens, M., Penaud, S., Rossignol, M., Cornet, F. and Boccard, F. 2004, Macrodomain organization of the Escherichia coli chromosome. EMBO J, 23, 4330–4341.

40. Boccard, F., Esnault, E. and Valens, M. 2005, Spatial arrangement and macrodomain organization of bacterial chromosomes. Molecular microbiology, 57, 9–16.

41. Dillon, S. C. and Dorman, C. J. 2010, Bacterial nucleoid-associated proteins, nucleoid structure and gene expression. Nat Rev Microbiol, 8, 185–195.

42. Monk, J. and Palsson, B. O. 2014, Genetics. Predicting microbial growth. Science, 344, 1448–1449.

