## Supplemental figures for "The common chromosomal periodicity of transcriptomes is correlated with the bacterial growth rate"

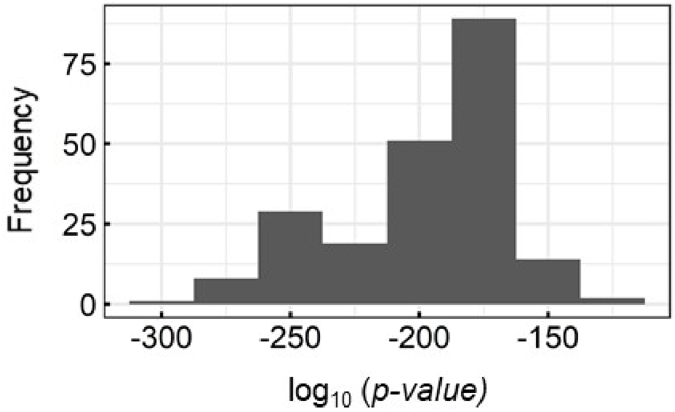


**Figure S1 Histogram of the significance of the chromosomal periodicity.** Fisher's g-test was performed to evaluate the statistical significance of the common six periods of the transcriptomes. The *p*-values are shown in the logarithmic scale.

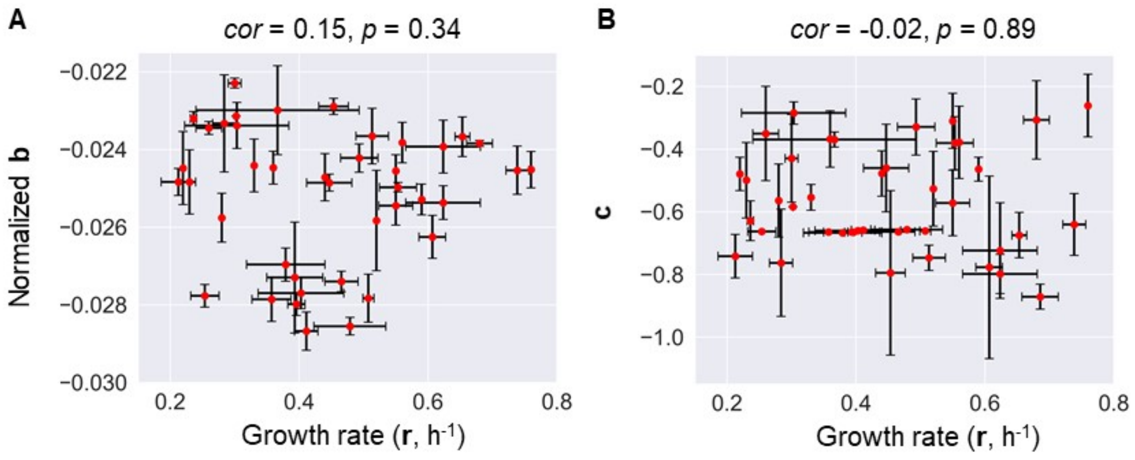


**Figure S2 Scatter plots of the periodic parameters (b and c) and the growth rate.** The standard errors of biological replicates are indicated. The correlation coefficient and its significance are indicated.

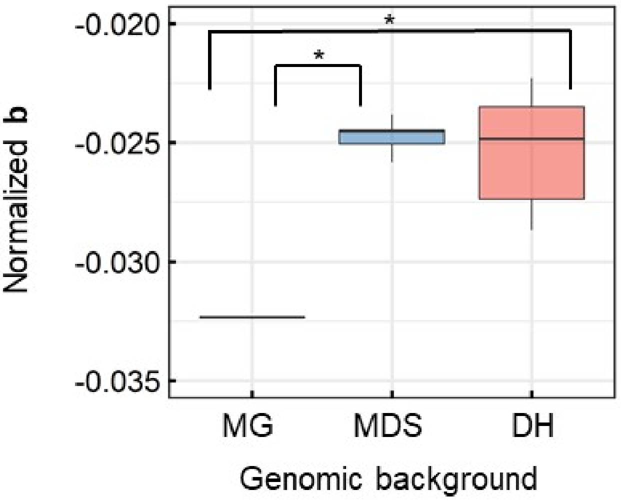


**Figure S3 Boxplot of the periodic parameter b.** The normalized parameter **b** was classified in accordance with the genomes. MG, MDS, and DH indicate the wild-type genome of MG1655, the partial genome of MDS42 and its derivatives, and the genomes of an assortment of genetically engineered DH1 strains, respectively. Asterisk indicates $p < 0.05$.

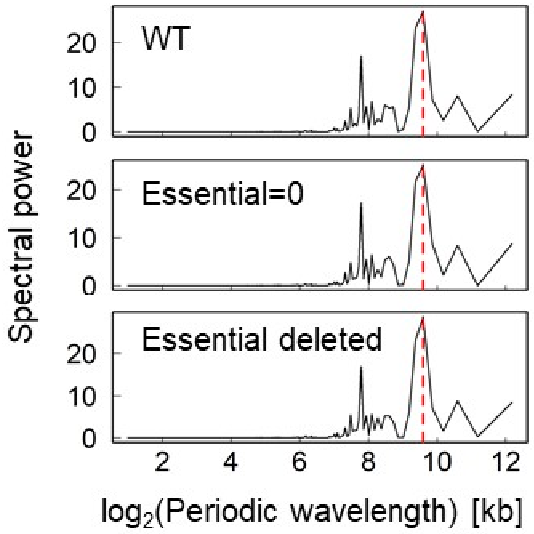


**Figure S4 Contribution of essential genes to chromosomal periodicity.** The distribution of the Fourier-transformed periodic wavelength at a logarithmic scale is shown. The MG1655 transcriptomes under normal conditions were subjected to analysis. The upper, middle and bottom panels represent the transcriptomes of all genes in the wild-type genome, which were determined by replacing the expression data of the essential genes with zero and removing the expression data of the essential genes. The most significant spectral power is indicated with a red broken line.

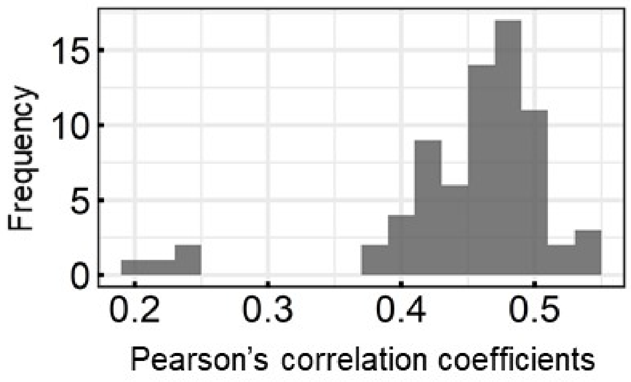


**Figure S5 Histogram of the correlation coefficients between GyrA binding activity and gene expression levels.** All transcriptomes were subjected to calculations, resulting in the 213 values that are shown in the histogram.

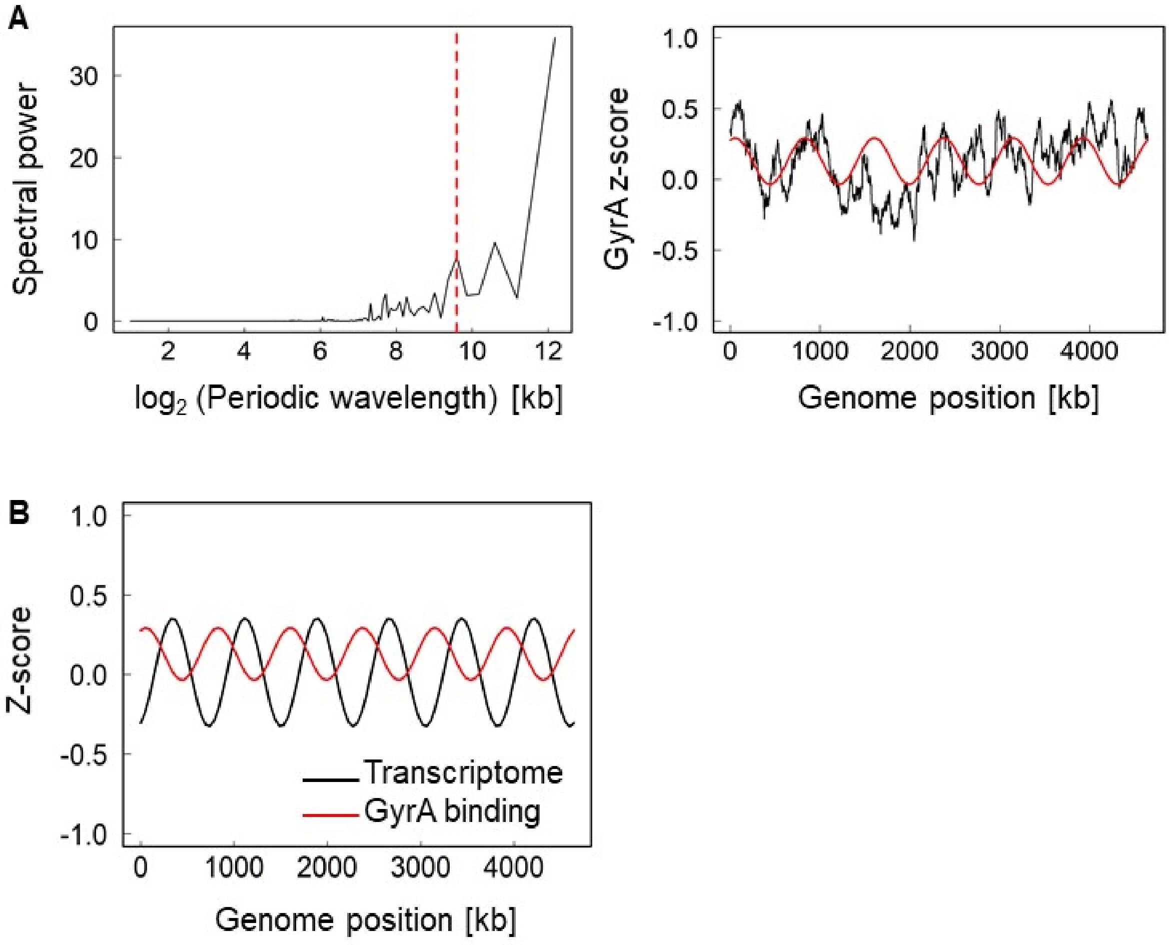


**Figure S6 Chromosomal periodicity of GyrA binding activity. A.** Fourier transform of the GyrA binding activity. The left and right panels represent the distributions of the Fourier-transformed periodic wavelengths at the logarithmic scale and the estimated chromosomal periodicity of the GyrA binding activity, respectively. The broken lines and solid curves in red indicate the power spectra corresponding to the six periods, as estimated by the Fourier transform and the corresponding fitted period of the GyrA binding activity, respectively. The GyrA binding activity (Chip-seq values) for every 1 kb sliding window and 100 kb smoothing are shown. **C.** Chromosomal periodicity of the transcriptome and GyrA binding activity. Red and black curves indicate the periodicity of the GyrA binding activity and the transcriptome, respectively. Both were calculated using a 1 kb sliding window and are shown as a 100 kb moving average.
